# IFNγ expression correlates with enhanced cytotoxicity in CD8^+^ T cells

**DOI:** 10.1101/2025.05.09.652839

**Authors:** Varsha Pattu, Elmar Kause, Hsin-Fang Chang, Jens Rettig, Xuemei Li

## Abstract

**Background:** CD8+ T lymphocytes (CTLs) act as serial killers of infected or malignant cells by releasing large amounts of interferon-gamma (IFNγ) and granzymes. Although IFNγ is a pleiotropic cytokine with diverse immunomodulatory functions, its precise spatiotemporal regulation and role in CTL-mediated cytotoxicity remain incompletely characterized.

**Methods:** Using wild-type (WT) and granzyme B-mTFP knock-in mice, we combined in vitro approaches, including T-cell isolation and culture, plate-bound anti-CD3e stimulation, degranulation assays, flow cytometry, immunofluorescence, and structured illumination microscopy, to investigate IFNγ dynamics in CTLs.

**Results:** IFNγ expression in CTLs was rapid, transient, and strictly dependent on T-cell receptor (TCR) activation. We identified two functionally distinct IFNγ-producing subsets: IFNγ^high^ (IFNγ^hi^) and IFNγ^low^ (IFNγ^lo^) CTLs. IFNγ^hi^ CTLs exhibited an effector/effector memory phenotype, significantly elevated CD107a surface expression (a marker of lytic granule exocytosis), and pronounced colocalization with cis-Golgi and granzyme B compared to IFNγ^lo^ CTLs. Furthermore, CRTAM, an early activation marker, correlated with IFNγ expression in naive CTLs.

**Conclusion:** Our findings establish a link between elevated IFNγ production and enhanced CTL cytotoxicity, implicating CRTAM as a potential regulator of early CTL activation and IFNγ induction. These insights provide a foundation for optimizing T cell-based immunotherapies against infections and cancers.

## Introduction

CD8^+^ cytotoxic T lymphocytes (CTLs) are critical immune sentinels that protect the host against infections and malignancies. Upon antigen recognition by specialized antigen-presenting cells (APCs), naive CD8^+^ T cells (T_N_) undergo clonal expansion and differentiate into short-lived effector CTLs (T_E_), which migrate to peripheral tissues and inflammatory sites^[1,2]^. Most T_E_ undergo apoptosis during the contraction phase^[3]^, while a small subset persists as long-lived memory T cells (T_M_). These memory populations consist of at least two distinct subsets: central memory (T_CM_) and effector memory (T_EM_) T cells^[4,5]^. In mice, short-lived T_E_ downregulate L-selectin (CD62L) but retain partial expression of IL-7 receptor alpha (CD127)^[6]^, whereas long-lived T_M_ constitutively express CD127. T_CM_ exhibit high CD62L expression, while T_EM_ downregulate CD62L. Notably, under chronic infections or cancer, persistent antigen exposure can drive naive T cells into a dysfunctional state termed T cell exhaustion (T_EX_)^[7]^. Understanding the heterogeneity and function of CTL substets is crucial to discern the complex dynamics of immune responses in various disease^[8]^.

CTLs eliminate target cells through lytic (granzymes and perforin) and non-lytic (IFNγ and TNFα) mechanisms^[9,10]^. Their effector functions are regulated by transcription factors such as T-bet, Eomes, CRTAM (Class-I MHC-restricted T-cell associated molecule), T-box, and Runx family proteins^[11–13]^. The killing capacity of CTLs depends on cytotoxic granules (CGs) and degranulation efficiency^[14]^. CGs are secretory lysosomes (SGs) containing granzymes and perforin^[15]^, enclosed by a lipid bilayer embedded with lysosome-associated membrane glycoproteins (LAMPs), including CD107a (LAMP-1), CD107b (LAMP-2), and CD63 (LAMP-3)^[16]^. Upon target recognition, CGs fuse with the plasma membrane, releasing their cytotoxic substances via exocytosis^[13]^. Since resting CTLs lack surface LAMPs, CD107a/b expression serves as a quantitative marker for degranulation^[12]^. Additionally, Fas ligand (FasL/CD95L), stored in specialized SGs, is externalized during degranulation to induce apoptosis in target cells^[17]^. Intriguingly, CTLs also express Fas receptor (Fas), enabling fratricidal killing (CTL-fratricide)—a mechanism that eliminates effector T cells during immune contraction^[17]^.

Beyond direct cytotoxicity, CTLs exert effector functions via interferon-gamma (IFNγ) and tumor necrosis factor-alpha (TNFα) secretion. IFNγ is primarily produced by NK cells, Th1 cells, and CD8^+^ T cells^[18,19]^. In murine models of *Mycobacterium tuberculosis* infection, CD4^+^ and CD8^+^ T cells are the dominant IFNγ producers in vivo^[20]^. IFNγ is a pleiotropic cytokine with dual roles—exhibiting antiviral^[21,22]^ and antitumor activity while paradoxically promoting tumor progression in certain contexts^[23–25]^. Mechanistically, IFNγ signaling upregulates interferon regulatory factor-1 (IRF-1), which enhances MHC-I/II expression, rendering target cells more susceptible to CTL attack^[26]^. As a proinflammatory cytokine, IFNγ also activates macrophages to secrete TNFα and IL-6^[27–29]^.

In this study, we investigated IFNγ production, subcellular localization, cellular subtypes, and cytotoxic function in IFNγ-expressing CTLs. We found that naive CD8^+^ T cells rapidly acquire IFNγ production within 2 hours of TCR activation whereas GzmB expression is restricted to effector CTLs with mature immune synapses^[30]^. Activated CTLs exhibited transient, TCR-dependent IFNγ expression, contrasting with pre-stored GzmB in resting CTLs. Strikingly, IFNγ^+^CTLs segregated into IFNγ^hi^ and IFNγ^lo^ subsets, while GzmB^+^CTLs were homogeneous. IFNγ^hi^ cells displayed a T_E_/T_EM_ phenotype (low CD62L), whereas IFNγ^lo^ cells resembled T_CM_ (high CD62L). Degranulation assays revealed that IFNγ^hi^ CTLs exhibited significantly higher surface CD107a than IFNγ^lo^ cells, suggesting superior lytic activity. Furthermore, IFNγ^hi^ CTLs showed pronounced colocalization with cis-Golgi and GzmB^+^ vesicles, reinforcing the link between IFNγ expression and cytotoxicity. Finally, comparative analysis of IFNγ and CRTAM expression in naive vs. activated CD8^+^ T cells suggested that CRTAM may regulate early T cell activation and IFNγ production.

## Methods

### Mice

Wild-type (WT) mice were purchased from Charles River, and granzyme B-mTFP knock-in (GzmB-mTFP KI) mice were generated as previously described^[30]^. All experimental procedures were conducted in compliance with regulations of the state of Saarland (Landesamt für Verbraucherschutz, AZ.: 2.4.1.1 and 11/2021).

### Cell isolation

Mice were anesthetized with CO_2_ and executed by cervical dislocation. The left abdominal cavity was exposed, and the spleen was carefully removed, placed on a 70 μm cell strainer (Corning Life Sciences) and ground. The grinding slurry remaining on the strainer was rinsed with RPMI medium, and the cell suspension (10 mL) was collected into a 15 mL sterile centrifuge tube (Corning Life Sciences) and centrifuged (6 min, 1100 rpm without a break) to wash the splenocytes. Washed splenocytes were mixed and incubated with erythrocyte lysis buffer (1 mL) for 30 seconds to lyse the erythrocytes therein. Immediately, 10 mL of ice-cold RPMI was added to terminate lysis and centrifuged (6 min, 1100 rpm), and the cell sediment was washed once more with isolation buffer to remove erythrocyte debris. Primary CD8^+^ T lymphocytes were positively isolated from spleens according to the instructions of the Dynabeads FlowComp Mouse CD8 Kit (Thermo Scientific).

### Cell culture

Isolated primary CD8^+^ T lymphocytes were cultured at a density of 1 × 10^6^ cells/mL in AIM V medium (Gibco) supplemented with 10% FCS, 50 μM β-Mercaptoethanol (BME), and 100 U/mL recombinant mouse IL-2 (Life technology) and stimulated with anti-CD3e/anti-CD28 magnetic beads (number of cells: magnetic beads = 1:0.8, Thermo Fisher Scientific) to generate activated CTLs. Cells were cultured in 24-well plates in a cell culture incubator (37 °C, 5% CO_2_, saturated humidity) according to the protocol of Bzeih et al. (2016)^[41]^. The first passaging of the cells was started after 2 days of culture and then once a day with the addition of fresh AIM V medium supplemented with 10% FCS, 50 μM BME and 100 U/mL recombinant IL-2. Usually, day3 to day5 cells were used for experiments.

For some experiments, cells were used immediately after the CD3e/CD28 activation beads had been removed. For most experiments cells were allowed to recover for 2 h after CD28/CD3e beads had been removed. Those cells underwent a “re-stimulation” during the experiment. By this is meant that by appropriate measures, the T-lymphocyte receptors have been acutely stimulated to trigger cytotoxic actions, such as the release of GzmB and IFNγ. For FACS experiments: To induce intracellular IFNγ-expression, day3-5 beads-activated WT CTLs were washed and restimulated with optimal concentration of plate-bound anti-CD3e antibody (10 μg/mL) from Estl et al. (2020)^[29]^ for 0, 0.5 h, 1 h, 2 h, 3 h, and 4 h, 6 h, 8 h, 12 h and 24 h in a cell culture incubator.

### Immunocytochemistry

CTLs were stained with an anti-IFNγ antibody to detect endogenous IFNγ. Therefore, cells were fixed with freshly prepared ice-cold 4% paraformaldehyde in DPBS (1×) and washed in DPBS (1×) containing 0.1 M glycine (quenching of autofluorescence). After a second wash cells were permeabilized with for 20 minutes in DPBS (1×) containing 0.1% Triton-X 100 (Roth), then blocked for 20-30 minutes in blocking buffer contained 5% BSA. Finally, cells were stained with primary rat anti-mouse IFNγ (1:200) and secondary anti-rat antibodies (1:1000), or rat anti-mouse IFNγ-Alexa488 (1:200), before they were mounted and imaged using high-resolution structured illumination microscopy (SIM).

Antinodies used in this study were listed in Extended Data Table 1.

### Structure illumination microscopy

#### Structured illumination microscopy setup

The SIM setup was from Zeiss (ELYRA PS.1). Images were acquired using a 63× Plan-Apochromat (NA1.4) objective with excitation light of 488, 561, and 647 nm and then processed for SIM to obtain higher resolutions. Z-stacks of 200 nm step size were used to scan cells. Zen 2012 software (Zen 2012; Carl Zeiss) was used for the acquisition and processing of the images for higher resolutions.

#### Colocalization analysis

For colocalization analysis the JACoP plugin (Bolte 2006) of Fiji (Johannes, 2012) was used. Pearson’s and Manders’ overlap coefficients (Manders et al., 1993) were used for quantification of the degree of colocalization.

### Flow cytometry

#### Intracellular IFNγ measurement

WT CTLs were washed with DPBS (1×) after restimulation for 0.5 h, 1 h, 2 h, 3 h, 4 h, 6 h, 8 h, 12 h, and 24 h with plate-bound anti-CD3e antibody (10 μg/mL). The pellet (0.5 × 10^6^ CTLs) was resuspended in 250 μL fixation buffer and incubated on ice for 10 min using BD Cytofix/ Cytoperm™ Fixation/Permeabilization Kit (554714, BD Bioscience). Next, CTLs were centrifuged, resuspended in 100 μL wash buffer (1×) and stained for intracellular IFNγ and GzmB with rat anti-mouse IFNγ-Alexa488 (1:200) and rat anti-mouse GzmB-Alexa647 (1:200)) on ice (1 h). Cells were washed and measured by flow cytometry (FACS, BD FACSAria III). Data were analyzed using FlowJo v10.10.0_CL software (Celeza-Switzerland).

#### Surface markers analysis

CD8^+^ T-lymphocytes either with or without (re)stimulation, were taken from cell culture and washed with cold-DPBS (1×). Next they were stained for CD62L, CD69 and CD25 using corresponding fluorescent Abs without fixation (live staining, 30 min on ice). Next, cells were washed twice with cold-wash buffer (1×) and resuspended in cold-DPBS (1×) for FACS analysis.

### Degranulation assay

Activated WT CTLs were washed and resuspended in AIM V after removing beads from the culture. Cells (0.2 × 10^6^/200 μL) were then cultured on a 10 µg/mL anti-CD3e coated or DPBS (1×) coated 96-well plate with 1 µL anti-CD107a-PE for 3 h at 37 °C for degranulation. The control cells were incubated without anti-CD107a-PE as a background signal. All cells were then washed twice with cold DPBS (1×) and checked by FACS. Data were analyzed using FlowJo software. Gates were set according to no fluorescence control cells. Each independent experiment had a duplicate.

### Statistical analysis

For data analysis and to calculate the statistical significance, Fiji, ImageJ-win64, Microsoft Excel (Microsoft), SigmaPlot 13 and GraphPad_Prism10.2.3 were used. The respective method to calculate statistical significance was given in the text for each figure. Figures were generated using FlowJo_v10.10.0_CL, GraphPad_Prism, Excel and PowerPoint.

## Results

### Transient IFNγ expression depends on TCR re-activation in resting CTLs

To determine whether interferon-gamma (IFNγ) expression is constitutive or regulated in activated CTLs, we analyzed intracellular IFNγ dynamics following TCR re-stimulation as previous report^[31]^. Day 3–5 activated CTLs, with or without re-stimulation, were fixed, permeabilized (BD Cytofix/Cytoperm™), and stained for IFNγ-Alexa488 and GzmB-Alexa647 (without Golgi transport inhibition). Flow cytometry data were acquired and analyzed using FlowJo v10.

IFNγ- and GzmB-expressing CTLs was shown in pseudocolor plots in 4 h TCR re-stimulation (Figure 1A,B). In unstimulated CTLs, IFNγ was nearly undetectable, but its expression increased rapidly upon TCR re-stimulation (Figure 1A). Whereas large amount of GzmB exisited before restimulation and had less variation after TCR re-stimulation (Figure 1B). The expression level of IFNγ and GzmB in CTLs exhibited a maturation-dependent pattern and distinct kinetics. IFNγ expression (measured by Median fluorescence intensity, MFI) showed a transient peak (Figure 1C), with day 5 CTLs displaying faster upregulation than day 3 CTLs at 2 h (*p* = 0.020), 3 h (*p* = 0.004), and 4 h (*p* = 0.004). Day 4 CTLs showed IFNγ MFI kinetics comparable to day 5 CTLs but significantly faster than day 3 at 3 h (*p* = 0.044). In contrast, GzmB MFI remained stable across day 3-5 CTLs during the 4-hour restimulation period (Figure 1D).

**Figure 1.**
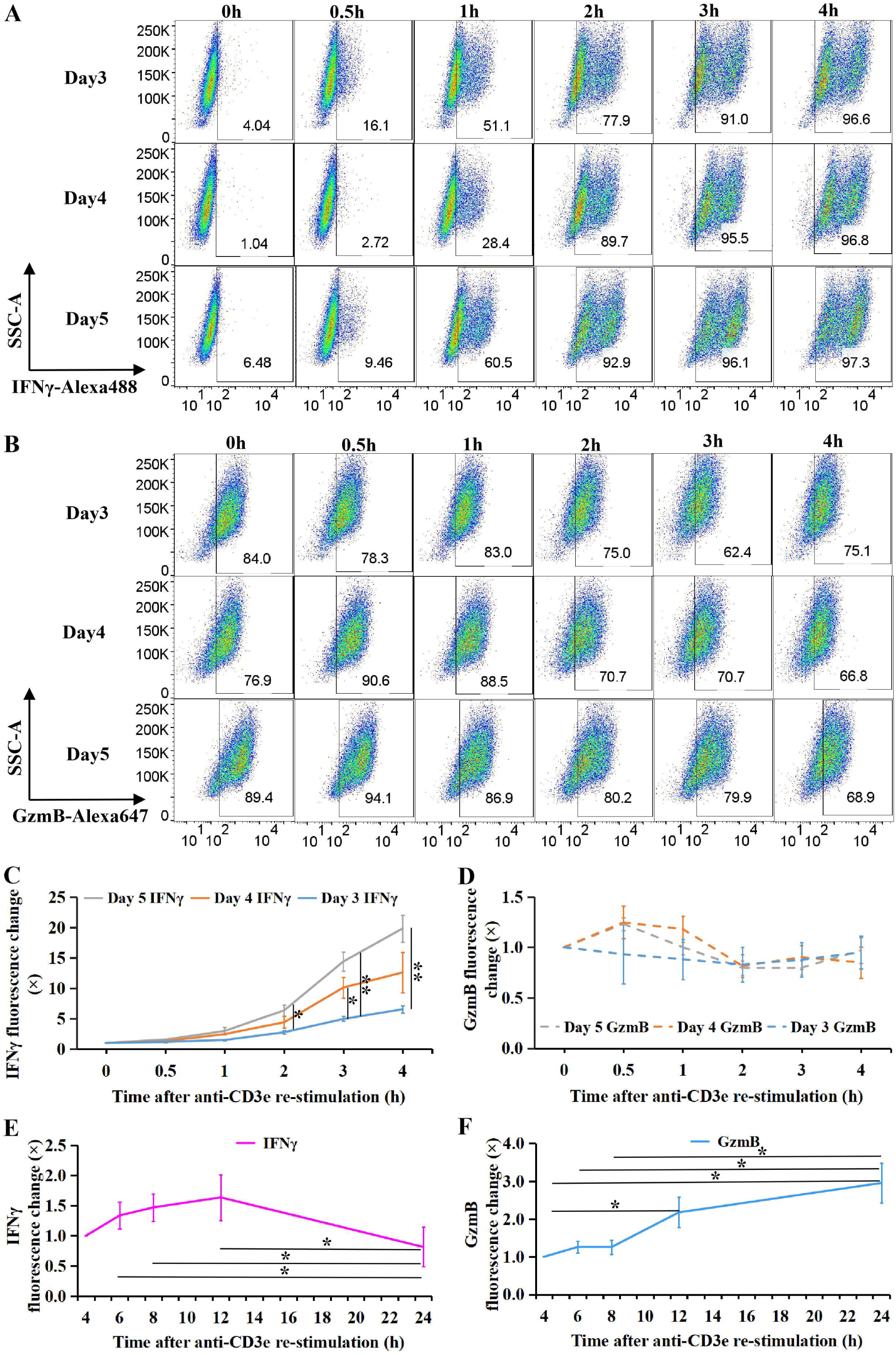
Intracellular IFNγ and granzyme B (GzmB) expression in activated CTLs. **(A–D)** WT CTLs (days 3–5 post-activation) were re-stimulated with plate-bound anti-CD3e antibody (10 μg/mL) and analyzed for intracellular IFNγ and GzmB expression over 0–4 h. **(A, B)** Pseudocolor plots depict fluorescence intensity of rat anti-mouse IFNγ-Alexa488 (x-axis) and GzmB-Alexa647 (x-axis) against side scatter area (SSC-A, y-axis). **(C, D)** Median fluorescence intensity (MFI) of IFNγ-Alexa488 and GzmB-Alexa647. **(E, F)** Temporal changes in IFNγ and GzmB expression in re-stimulated CTLs. MFI dynamics of IFNγ-Alexa488 and GzmB-Alexa647. Data are mean ± SEM (N ≥ 3). Statistical significance was determined by one-way ANOVA and unpaired t-test (\**p* < 0.05, \*\**p* < 0.01, \*\*\**p* < 0.001; NS, not significant).

Longitudinal analysis revealed divergent temporal patterns: IFNγ levels peaked at 12 h before declining sharply by 24 h, while GzmB accumulated continuously. IFNγ MFI was significantly elevated at 6 h, 8 h, and 12 h compared to 24 h (*p* = 0.021, *p* = 0.017, and *p* = 0.021, respectively) (Figure 1E). Conversely, GzmB MFI showed progressive accumulation, with significantly higher levels at 24 h versus 4 h, 6 h, and 8 h (*p* = 0.021, *p* = 0.037, and *p* = 0.039), and at 12 h versus 4 h (*p* = 0.042) (Figure 1F).

In conclusion, in activated CTLs, the expression of IFNγ and GzmB exhibited a maturation-dependent pattern and distinct kinetics. IFNγ expression is transient, but GzmB expression lasts longger compared to IFNγ after TCR (re-)activation.

### CTL subtype dynamics upon TCR re-stimulation

Cytotoxic T lymphocytes (CTLs) comprise distinct functional subsets, including effector cells (T_E_), effector-memory cells (T_EM_), and central-memory cells (T_CM_). To determine whether IFNγ expression is subtype-specific and whether TCR re-stimulation alters the subtype composition of cultured CTLs, we stimulated activated CTLs with 10 μg/mL plate-bound anti-CD3e^[31]^ for 0.5–4 h and analyzed subtype markers (CD44^hi^ for T_E_/T_EM_, CD62^hi^ for T_CM_) by flow cytometry at multiple timepoints.

Neither the percentage of CD44^+^ cells (Extended Data Figure 1A) nor CD44 MFI (Extended Data Figure 1B) changed significantly during 4 h re-stimulation, regardless of culture day (day 3–5), indicating that overall effector differentiation status of the culture remained stable, independent of TCR re-stimulation or culture duration. In contrast, CD62L^+^ cell frequency (Extended Data Figure 1C) and CD62L MFI (Extended Data Figure 1D) declined sharply upon re-stimulation (*p* < 0.05), indicating a transition from T_CM_ to T_EM_ (Figure 2A–C).

**Figure 2.**
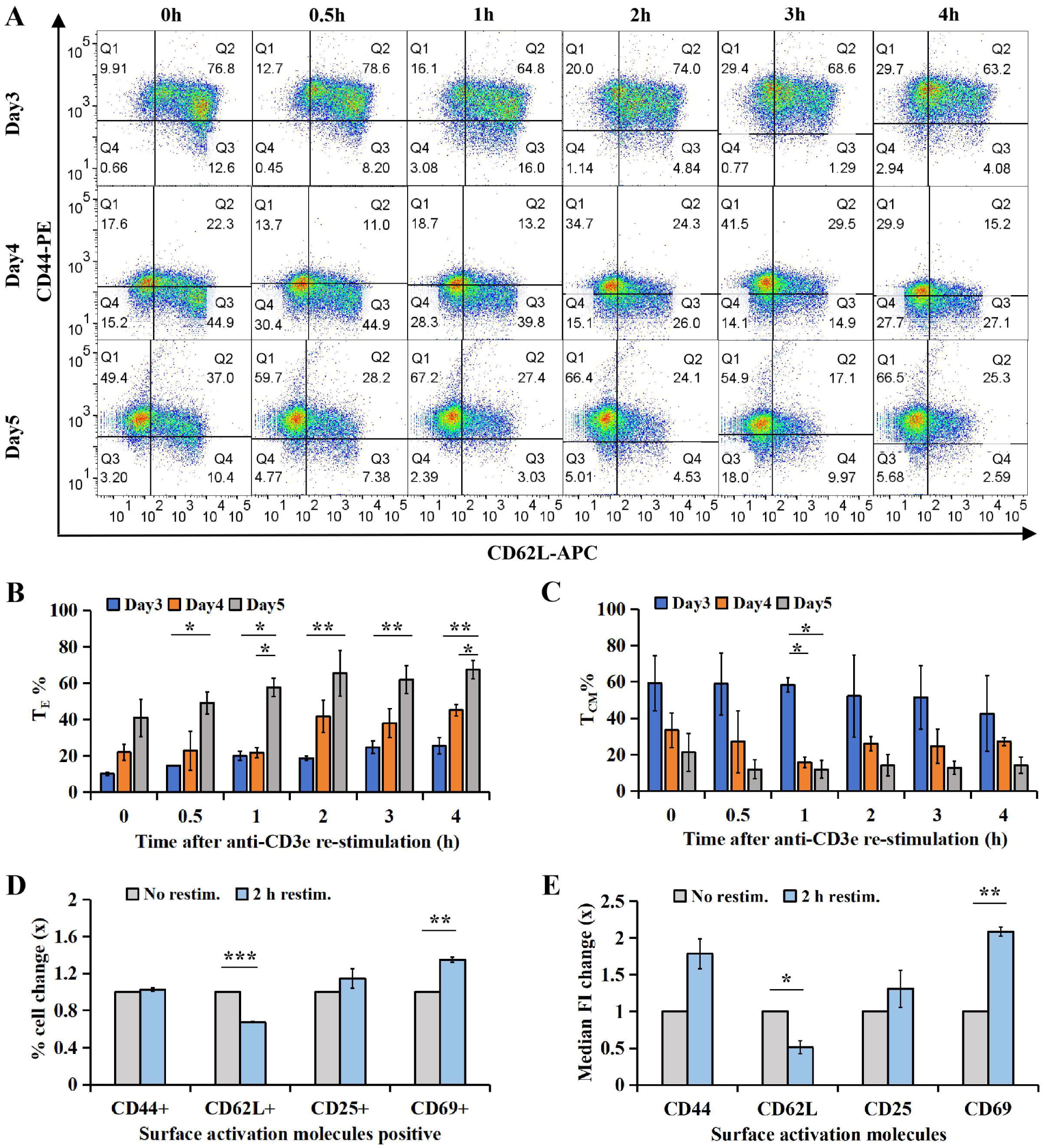
Surface activation marker expression profiles in CTLs analyzed by flow cytometry. **(A–C)** WT CTLs (days 3–5 post-activation) were re-stimulated with plate-bound anti-CD3e antibody (10 μg/mL) for 0–4 h and stained for CD44 and CD62L. **(A)** Pseudocolor plots depict fluorescence intensity of rat anti-mouse CD62L-APC (x-axis) and CD44-PE (y-axis). **(B, C)** Frequencies of T_E_ and T_CM_ populations. **(D, E)** WT CTLs (day 5 post-activation) were re-stimulated with plate-bound anti-CD3e (10 μg/mL) for 2 h and analyzed for multiple activation markers. **(D)** Fold change in CD44+, CD62L+, CD25+, and CD69+ cell populations. **(E)** MFI change of CD44-PE, CD62L-APC, CD25-PE, and CD69-APC before and after 2 h restimulation. Data represent mean ± SEM (N ≥ 3). Statistical analysis was performed using one-way ANOVA and unpaired t-test (\**p* < 0.05, \*\**p* < 0.01, \*\*\**p* < 0.001; NS, not significant).

Pseudocolor plots in quadrant (Figure 2A) revealed subtype distribution. Cells in Q1 (CD44+CD62-) were T_E/EM_, Q2 (CD44+CD62+) were T_CM_, Q3 (CD44-CD62+) were T_N._ Day 5 CTLs exhibited significantly more T_E_/T_EM_ than day 3 CTLs at 4 h and day 4 CTLs at 1 h/4 h (*p* < 0.05; Figure 2B). Concurrently, T_CM_ frequency was lower in day 5 vs. day 3/day 4 CTLs at 1 h (*p* < 0.05; Figure 2C), confirming that longer culture promotes T_E/EM_ differentiation upon re-stimulation.

Screening of activation markers (CD25, CD44, CD62L, CD69) in day 5 CTLs revealed robustly decreased CD62L and increased CD69 in both cell frequency and MFI (Figure 2D,E) after 2 h re-stimulation, while no significant changes were found in CD44 or CD25, implicating CD62L downregulation and CD69 upregulation are robust indicators of TCR-driven T_CM_-to-T_EM_ transition.

### Cellular subtype and cytotoxicity analysis of IFNγ^hi^ and IFNγ^lo^ CTLs

To determine whether IFNγ expression correlates with cytotoxic T lymphocyte (CTL) effector function, we analyzed reactivated CTLs, which exhibited distinct clusters of high (IFNγ^hi^) and low (IFNγ^lo^) IFNγ expression (Figure 1A and Figure 3A, row 1). Using a fluorescence intensity threshold of 10³, we classified IFNγ-positive CTLs into IFNγ^hi^ and IFNγ^lo^ subsets (Figure 3A, row 2). Longed re-stimulation led to increased effector T cell (T_E_) generation (Figure 2B & 3A row3). IFNγ^hi^ subset generated significantly higher ratio of T_E_ than IFNγ^lo^ subset at 3 and 4 hours of restimulation (Figure 3B).

**Figure 3.**
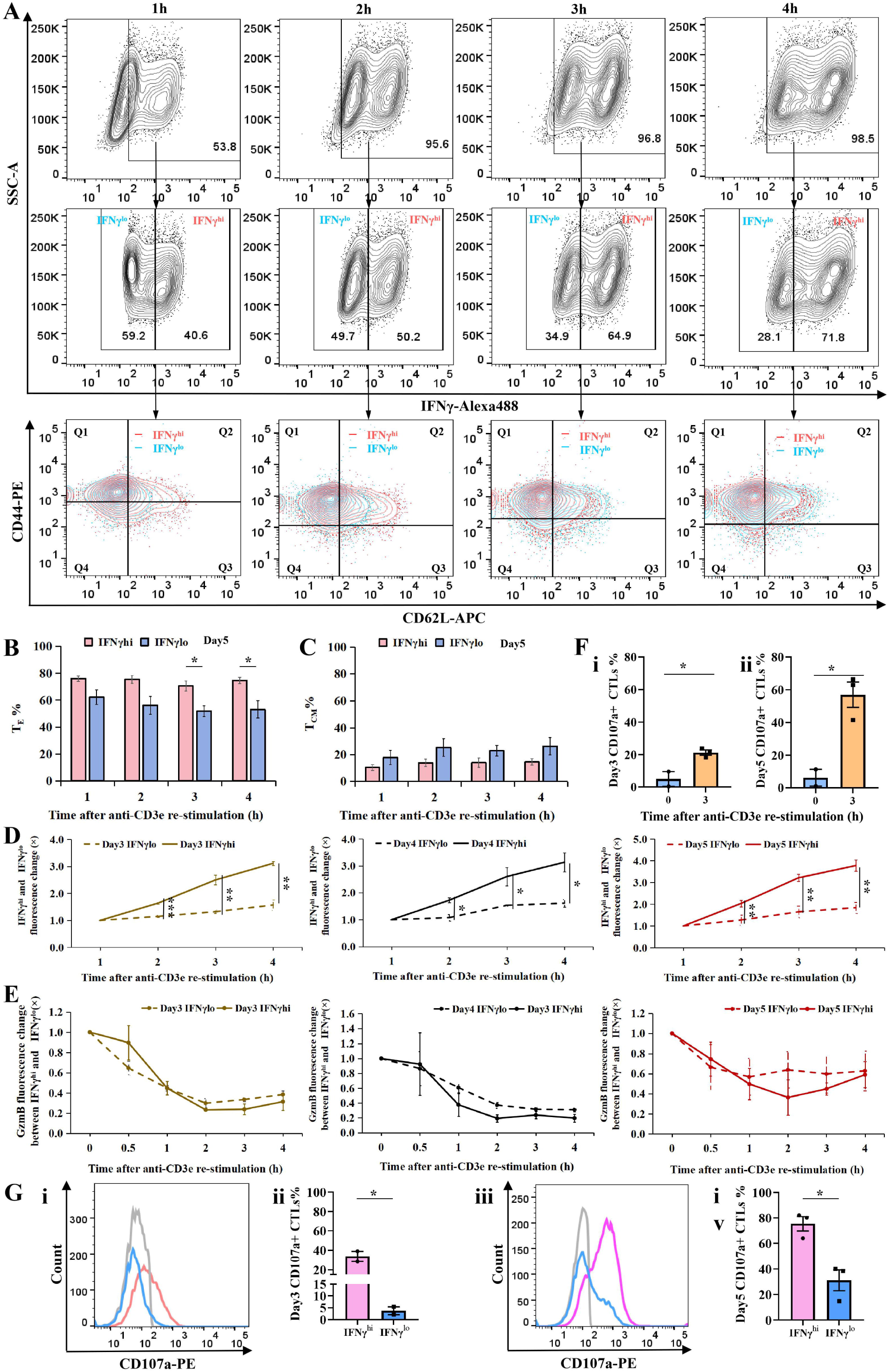
Characterization of IFNγ^hi^ and IFNγ^lo^ CTL subsets and their cytotoxic potential. **(A-E)** WT CTLs were re-stimulated with plate-bound anti-CD3e antibody for 0-4 h and processed for flow cytometry analysis: One aliquot was surface-stained with anti-CD44-APC and anti-CD62L-PE, followed by intracellular staining for IFNγ-Alexa488. Another aliquot was intracellularly stained for IFNγ-Alexa488 and GzmB-Alexa647 **(A)** Gating strategy and surface marker expression profiles of IFNγ^hi^ vs IFNγ^lo^ CTLs. **(B)** Percentage of effector T cells (T_E_) in each subset. **(C)** Percentage of central memory T cells (T_CM_) in each subset. **(D)** IFNγ fluorescence intensity in IFNγ^hi^ vs IFNγ^lo^ subsets. **(E)** GzmB fluorescence intensity in IFNγ^hi^ vs IFNγ^lo^ subsets. **(F,G)** Degranulation capacity assessed by CD107a surface expression: Day 3 and 5 WT CTLs were stimulated with plate-bound anti-CD3e for 3 h in the presence of anti-CD107a-PE, followed by intracellular IFNγ staining. **(F)** Percentage of CD107a+ CTLs pre- and post-stimulation. **(G) i,iii**: CD107a expression in IFNγ^hi^ vs IFNγ^lo^ CTLs at day 3 and 5. **ii,iv**: CD107a fluorescence intensity in IFNγ^hi^ vs IFNγ^lo^ CTLs at day 3 and 5. Data represent mean ± SEM (N≥2 independent experiments). Statistical significance was determined by unpaired t-test (\**p*<0.05, \*\**p*<0.01, \*\*\**p*<0.001).

Although IFNγ^hi^ and IFNγ^lo^ CTLs showed comparable CD44 expression, IFNγ^hi^ CTLs exhibited significantly lower CD62L levels compared to IFNγ^lo^ CTLs (Supplement 2). The proportion of T_CM_ was slightly higher in IFNγ^lo^ CTLs, though not statistically significant (Figure 3C). When assessing IFNγ production kinetics, IFNγ^hi^ CTLs displayed markedly higher IFNγ MFI than IFNγ^lo^ CTLs at 2, 3, and 4 hours post-stimulation (Figure 3D), whereas GzmB MFI did not differ significantly between the two subsets (Figure 3E).

Extended TCR stimulation revealed a pronounced divergence in IFNγ fluorescence between IFNγ^hi^ and IFNγ^lo^ CTLs by 24 hours (*p* = 0.001), while GzmB fluorescence remained comparable (Extended Data Figure 3). In IFNγ^hi^ CTLs, IFNγ fluorescence peaked at 12 hours before declining significantly by 24 hours (vs. 4 h, *p* = 0.042 and *p* = 0.0025). In contrast, IFNγ^lo^ CTLs exhibited a gradual increase, peaking at 12 hours without subsequent decline, and showed higher fluorescence at 6, 12, and 24 hours than at 4 hours (*p* = 0.0483, *p* = 0.0112, and *p* = 0.0063, Extended Data Figure 3A). GzmB fluorescence in IFNγ^hi^ CTLs rose significantly at 6, 8, and 24 hours (vs. 4 h, *p* = 0.0022, *p* = 0.0075, and *p* = 0.0213), whereas in IFNγ^lo^ CTLs showed increased GzmB only at 12 and 24 hours (vs. 4 h, *p* = 0.0246 and *p* = 0.0171, Extended Data Figure 3B).

To evaluate cytotoxic capacity, we measured CD107a surface expression, a marker of degranulation. Unstimulated CTLs exhibited minimal CD107a expression (∼5%), but upon 3-hour re-stimulation, levels surged to 21.33% (day 3) and 57% (day 5), with day 5 showing significantly higher degranulation than day 3 (*p* = 0.0109; Figure 3F), suggesting that degranulation capacity escalates with CTL maturation. Notably, IFNγ^hi^ CTLs consistently displayed higher CD107a expression than IFNγ^lo^ CTLs (Figure 3G). Flow cytometry histograms depicted CD107a fluorescence intensity in unlabeled (gray), IFNγ^hi^ (pink), and IFNγ^lo^ (blue) CTLs (Figures 3Gi and 3Giii). On day 3, 33.75% of IFNγ^hi^ CTLs underwent degranulation versus only 3.76% of IFNγ^lo^ CTLs (Figure 3Gii). By day 5, degranulation frequencies rose to 75.47% (IFNγ^hi^) and 31.13% (IFNγ^lo^) (Figure 3Giv). These results suggests that cytotoxic activity correlates positively with IFNγ expression and cellular maturation, underscoring the functional superiority of IFNγ^hi^ CTLs.

### Subcellular localization analysis of IFNγ in IFNγ^hi^ and IFNγ^lo^ CTLs

To characterize the spatial distribution of IFNγ in cytotoxic T lymphocytes (CTLs), we analyzed the Pearson’s correlation coefficient (PCC) between endogenous IFNγ, cis-Golgi, and granzyme B (GzmB) in IFNγ^hi^ and IFNγ^lo^ subsets.

#### IFNγ colocalized with the Golgi Apparatus

Wild-type (WT) CTLs were re-stimulated with plate-bound aCD3e and stained for endogenous IFNγ and cis-Golgi. Super-resolution structured illumination microscopy (SIM) was performed using ZEN software, and PCC was quantified with Fiji. IFNγ exhibited extensive colocalization with cis-Golgi in WT CTLs. No significant difference in IFNγ-Golgi PCC was observed between IFNγ^hi^ and IFNγ^lo^ CTLs at 2 hours post-stimulation on day 4 (Figure 4A, B), while significantly higher colocalization in IFNγ^hi^ CTLs was displayed than IFNγ^lo^ CTLs after 4 hours of re-stimulation on day 2 (Figure 4C, D).

**Figure 4.**
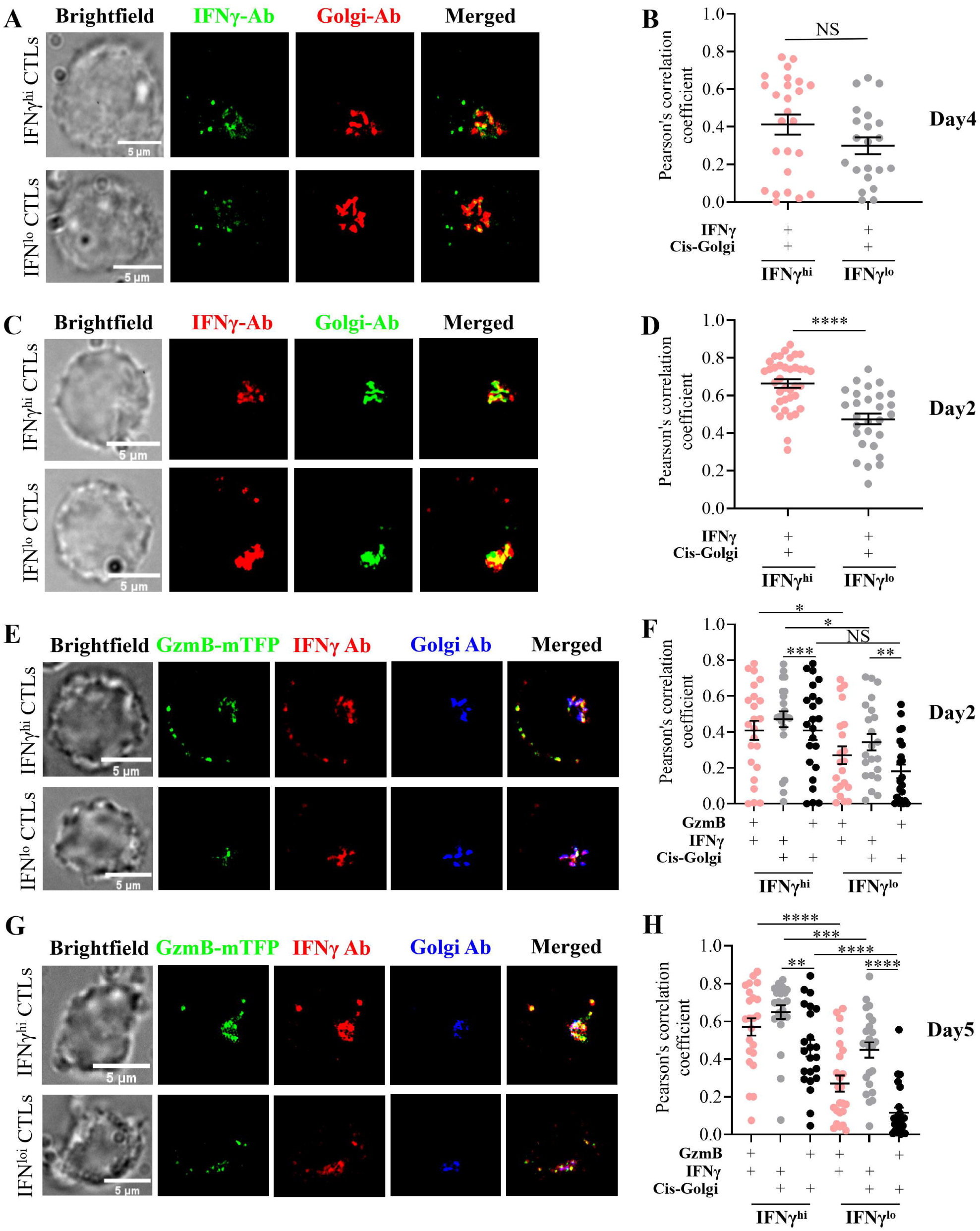
Subcellular localization of IFNγ in IFNγ^hi^ and IFNγ^lo^ CTL subsets. **(A-H)** WT or GzmB-mTFP KI CTLs were re-stimulated with plate-bound anti-CD3e (10 μg/mL) (for day2: 4 h restimulation, day4 and day 5: 2 h restimulation) and processed for super-resolution microscopy (SIM): Imaging and analysis: Cells were stained for IFNγ, GzmB, and cis-Golgi marker (GM130) IFNγ^hi^ vs IFNγ^lo^ subsets were imaged using optimized laser power and EMCCD gain settings. Colocalization analysis performed using Fiji-win64 software. Quantitative comparisons and graphing performed using GraphPad Prism 10.2.3. Experimental details: **(A-B)** Day 4 WT CTLs: Primary antibodies: Rat anti-mouse IFNγ-Alexa488 (1:200), mouse anti-GM130 (1:100). Secondary antibody: Goat anti-mouse-Alexa647 (1:1000). **(B)** Pearson’s correlation coefficient for IFNγ and cis-Golgi colocalization. **(C-D)** Day 2 WT CTLs: Primary antibodies: Rat anti-mouse IFNγ (1:200), mouse anti-GM130 (1:100). Secondary antibodies: Alexa647 chicken anti-rat IgG (1:800), Alexa568 goat anti-mouse IgG (1:400). Imaging parameters: IFNγ^lo^: 561 nm laser (1.5%, gain 80); 647 nm laser (1.5%, gain 50). IFNγ^hi^: 561 nm laser (1.2%, gain 80); 647 nm laser (0.6%, gain 25). **(D)** Pearson’s correlation coefficient analysis. **(E-H)** GzmB-mTFP KI CTLs: Primary antibodies: Rat anti-mouse IFNγ (1:200), mouse anti-GM130 (1:100). Secondary antibodies: Alexa647 chicken anti-rat IgG (1:800), Alexa568 goat anti-mouse IgG (1:400). **(E,F)** Day 2 CTLs imaging parameters: IFNγ^hi^: 488 nm (22%, gain 80); 568 nm (1.2%, gain 80); 647 nm (0.7%, gain 40). IFNγ^lo^: 488 nm (26%, gain 80); 568 nm (1.2%, gain 80); 647 nm (1.5%, gain 80). **(G,H)** Day 5 CTLs imaging parameters: IFNγ^hi^: 488 nm (18%, gain 80); 561 nm (1.2%, gain 80); 647 nm (0.3%, gain 20). IFNγ^lo^: 488 nm (16%, gain 80); 561 nm (1.5%, gain 80); 647 nm (1.0%, gain 80). **(F,H)** Pearson’s correlation coefficient analyses. Statistical analysis: Data represent mean ± SEM. Significance was determined by one-way ANOVA with t-tests (\**p*<0.05, \*\**p*<0.01, \*\*\**p*<0.001; NS, not significant).

#### IFNγ localized in cytotoxic granules (CGs)

To determine whether IFNγ traffics to CGs, we analyzed IFNγ and GzmB colocalization in GzmB-mTFP knock-in (KI) CTLs on days 2 and 5 post-stimulation. GzmB is a marker for CGs. SIM imaging (Figure 4E, F) and Fiji-based quantification (Figure 4G, H) revealed that a subset of IFNγ localized in GzmB+ CGs, with IFNγ^hi^ CTLs exhibiting significantly greater IFNγ-GzmB association than IFNγ^lo^ CTLs (Figure 4F, H).

#### Comparative localization in Golgi vs. CGs

IFNγ showed robust cis-Golgi localization in GzmB-mTFP KI CTLs, with IFNγ^hi^ CTLs displaying higher enrichment than IFNγ^lo^ CTLs (Figure 4F, H). In contrast, GzmB exhibited minimal Golgi association, though IFNγ^hi^ CTLs on day 5 demonstrated slightly higher GzmB-Golgi PCC than IFNγ^lo^ CTLs (Figure 4H). No such difference was observed on day 2 (Figure 4F).

These findings demonstrate that: The IFNγ^hi^ subset retains higher IFNγ expression and greater IFNγ localization within CGs compared to IFNγ^lo^ CTLs. Newly synthesized IFNγ accumulates more prominently in the Golgi apparatus than GzmB following acute TCR stimulation. The preferential trafficking of IFNγ to CGs in IFNγ^hi^ CTLs suggests a direct role in cytotoxicity, consistent with their enhanced effector function. IFNγ expression is dynamically regulated, with transient Golgi-associated production preceding effector granule loading. Together, these data support a model wherein IFNγ^hi^ CTLs achieve superior cytotoxic activity through coordinated IFNγ synthesis, Golgi processing, and granule loading.

### Temporal regulation of CRTAM and IFNγ in naive and activated CTLs

To investigate the relationship between activation markers and IFNγ expression during early and late T cell activation, we analyzed IFNγ, CRTAM, CD62L, CD69, and CD25 expression in wild-type CD8^+^ T cells at day 0 (naive) and day 5 (activated) following TCR stimulation or restimulation.

#### Kinetics of CRTAM and IFNγ expression

Both CRTAM and IFNγ were upregulated upon TCR stimulation (Figure 5A-F, Extended Figure 4A-C). CRTAM^+^CTLs increased from 5.93% (1 h) to a peak of 96.6% (24 h) before declining to 10.4% by 72 h, while IFNγ^+^CTLs rose from 5.24% (1 h) to 35.6% (24 h) and subsequently decreased to 9.62% (72 h) (Extended Figure 4B). MFI for both molecules peaked at 24 h and became nearly undetectable by 72 h (Extended Figure 4C). Notably, day 2 and day 3 CTLs regained CRTAM and IFNγ expression upon restimulation (4 h and 3 h, respectively), confirming their transient, activation-dependent regulation (Extended Figure 4A).

**Figure 5.**
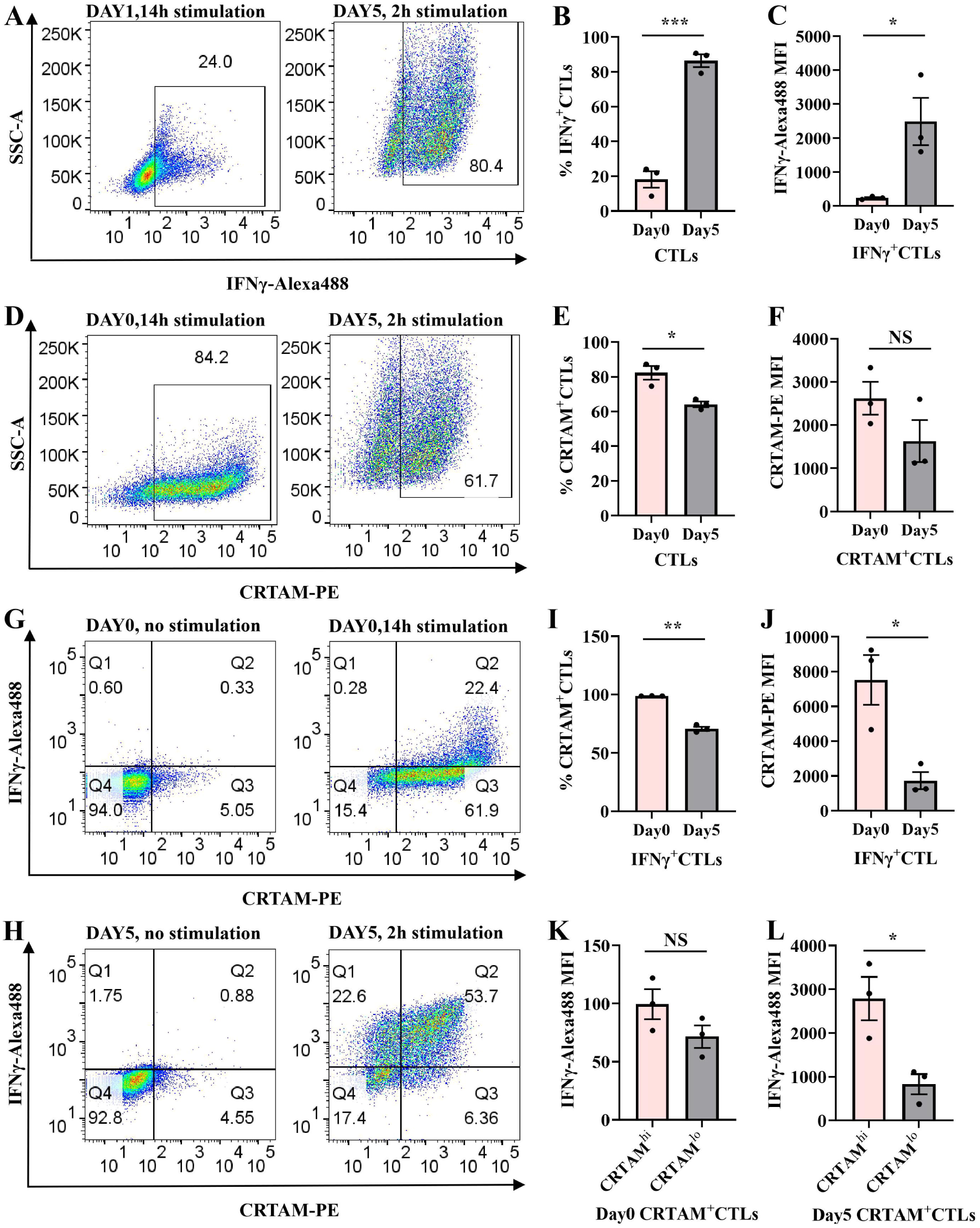
Co-expression analysis of CRTAM and IFNγ in activated CTLs by flow cytometry. Experimental design: Day 0 CTLs: Stimulated with plate-bound anti-CD3e (10 μg/mL) and anti-CD28 (5 μg/mL) for 14 h. Day 5 CTLs: Re-stimulated with plate-bound anti-CD3e (10 μg/mL) for 2 h in 96-well plates. Staining: Anti-CRTAM-PE and anti-IFNγ-Alexa488 antibodies. Flow cytometry analysis: **(A)** Representative pseudocolor plots of IFNγ expression in day 0 vs day 5 CTLs. **(B)** Frequency of IFNγ^+^CTLs. **(C)** MFI of IFNγ-Alexa488. **(D)** Representative pseudocolor plots of CRTAM expression. **(E)** Frequency of CRTAM^+^CTL populations. **(F)** MFI of CRTAM-PE. Co-expression analysis: **(G, H)** Quadrant plots showing IFNγ and CRTAM co-expression patterns. **(I)** Frequency of CRTAM^+^ cells in IFNγ^+^ CTLs. **(J)** MFI of CRTAM-PE in IFNγ^+^CTLs. **(K-L)** MFI of IFNγ-Alexa488 in CRTAM^hi^ vs CRTAM^lo^ subsets. Statistical analysis: Data represent mean ± SEM (N=3 independent experiments). Significance was determined by unpaired t-test (\**p*<0.05, \*\**p*<0.01, \*\*\**p*<0.001; NS, not significant).

#### Comparative expression of CRTAM and IFNγ in naive vs. activated CTLs

Day 5 CTLs exhibited significantly higher percentage of IFNγ^+^CTLs and IFNγ MFI than day 0 CTLs (Figure 5A-C), whereas naive CTLs had a greater proportion of CRTAM^+^CTLs (Figure 5D,E). CRTAM MFI, however, did not differ significantly between day 0 and day 5 (Figure 5F). Quadrant analysis of pseudocolor plots revealed distinct coexpression patterns.

In day 0 CTLs, only 20–25% of CRTAM^+^CTLs expressed IFNγ, but nearly 100% of IFNγ^+^CTLs were CRTAM^+^ after 14 h stimulation (Figure 5G, I; Extended Figure 4A). In day 5 CTLs, ∼71% of IFNγ^+^CTLs coexpressed CRTAM (Figure 5H, I).

Naive IFNγ^+^CTLs also displayed higher CRTAM MFI than their day 5 counterparts (Figure 5J). Strikingly, CRTAM^hi^ CTLs produced more IFNγ than CRTAM^lo^ subsets (Figure 5K, L), underscoring a positive correlation between CRTAM and IFNγ.

#### Activation marker dynamics

CRTAM upregulation in naive CTLs occurred as rapidly as CD69 induction, preceding CD25 expression (1–6 h; Extended Figure 4A,D,G). Conversely, CD62L levels declined post-stimulation (Extended Figure 4J). By day 2, CTLs entered contraction/memory phases, marked by rising CD62L and declining CD69/CD25 (Extended Figure 4E,F,I,L). With TCR restimulation, day2/3 CTLs were marked with reducing CD62L and increasing CD69/CD25 (Extended Figure 4E,F,H,I,L), while the percentage of CD25 didn’t change (Extended Figure 4K).

Our findings demonstrate that CRTAM and IFNγ are transiently expressed in a TCR-dependent manner. CRTAM serves as an early activation marker and correlates with IFNγ production in naive CTLs. CRTAM, CD62L, CD69, and CD25 collectively define CTLs’ activation states. These results position CRTAM as a key early responder during primary CTLs’ activation, functionally linked to IFNγ-mediated effector responses.

## Discussion

Effector CD8^+^ T cells (CTLs) are central to antimicrobial and antitumor immunity, employing both lytic and non-lytic cytotoxic mechanisms. While their lytic function mediated by GzmB and perforin is well-characterized^[30,32]^, the role of interferon-gamma (IFNγ) in CTL-mediated cytotoxicity remains less defined. Our study elucidates the dynamic expression profile of IFNγ in activated CTLs and its functional contribution to their cytotoxic potential.

### IFNγ dynamics and cytotoxic function

We demonstrate that IFNγ expression in CTLs is rapidly induced upon TCR activation, peaking early before declining transiently compared to the sustained expression of GzmB (Figure 1A–G). This transient expression coincided with CTL maturation, as evidenced by the progressive shift toward effector T cell (T_E_) phenotypes and the downregulation of CD62L (Figure 2). Notably, CTLs segregated into distinct IFNγ^hi^ and IFNγ^lo^ subsets, with the former exhibiting a stronger effector phenotype (Figure 3A–C).

Intriguingly, IFNγ^hi^ CTLs displayed significantly enhanced degranulation capacity (Figure 3D,G) despite comparable GzmB levels to IFNγ^lo^ CTLs (Figure 3E), suggesting that IFNγ potentiates lytic cytotoxicity independently of intracellular GzmB expression level. This was further supported by super-resolution imaging, which revealed greater colocalization of IFNγ with GzmB-positive cytotoxic granules in IFNγ^hi^ CTLs (Figure 4E–H). These findings position IFNγ not only as an immunomodulatory cytokine but also as a direct contributor to the lytic efficiency of CTLs. This finding is aligned with our previous research demonstrating that lytic IFNγ is stored within GzmB-containing cytotoxic granules and co-secreted at the immune synapse by effector CD8^+^ T cells, a mechanism that enhances cytotoxic T lymphocytes’ (CTLs) killing capacity^[33]^. Bhat et al. (2017) demonstrated that autocrine IFNγ production by CTLs enhances their motility and facilitates killing of primary target keratinocytes, both in vitro and in vivo. This finding highlights the critical dependence of CD8^+^ T cell cytotoxic function on local IFNγ signaling, which has important implications for immunotherapy targeting chronic viral infections and cancers^[34]^.

### CRTAM as a regulator of early CTL activation

The mechanisms governing IFNγ production in CTLs remain incompletely understood. Our data highlight CRTAM as a key early activation marker that correlates with IFNγ expression. CRTAM was upregulated transiently in naive CTLs upon TCR stimulation, preceding IFNγ induction (Figure 5; Extended Data Figure 3). Nearly all IFNγ^+^CTLs coexpressed CRTAM during primary activation (Extended Data Figure 3A), and CRTAM^hi^ subsets produced significantly more IFNγ than CRTAM^lo^ cells (Figure 5K,L).

The role of CRTAM in CTL biology aligns with its known functions in promoting IFNγ production in CD8^+^ T cells and CD4^+^ T cells^[35,36]^, in regulating CD4^+^ T cell polarization and in determing CD4^+^ cytotoxic T lymphocyte lineage^[35,37]^. Its interaction with CADM1 may facilitate early CTL priming by retaining activated cells in lymphoid tissues^[11]^, while its decline in day 5 CTLs (Figure 5I,J) suggests a stage-specific role in bridging early activation with effector differentiation.

### Future directions

While our work establishes a link between IFNγ expression and CTL cytotoxicity, the molecular pathways connecting CRTAM to IFNγ regulation warrant further investigation. Single-cell RNA sequencing could delineate the transcriptional programs distinguishing IFNγ^hi^ and IFNγ^lo^ subsets^[38–40]^, offering insights into their functional specialization. Additionally, mechanistic studies are needed to clarify whether IFNγ enhances lytic function through granule trafficking and fusion, target cell sensitization, or other effector pathways.

## Conclusions

In conclusion, the expression of IFNγ and GzmB shows a maturation-dependent pattern but dfifferent kinetics. Unlike GzmB, IFNγ production is rapid and transient following TCR (re)activation and does not require mature immune synapse formation. We further demonstrate that CD62L downregulation and CD69 upregulation serve as robust markers of TCR-driven transition from central memory (T_CM_) to effector memory (T_EM_) phenotypes. Functionally, IFNγ may enhance CTL cytotoxicity by promoting Golgi-dependent processing and efficient sorting into lytic granules, with elevated IFNγ levels correlating with greater killing capacity. Additionally, we identify CRTAM as a critical early responder in primary CTL activation, where it may regulate initial IFNγ expression and contribute to IFNγ-mediated effector functions. Collectively, these insights establish a mechanistic framework for refining T cell-based immunotherapies against infectious diseases and malignancies.

## Abbreviations

CTL: CD8+ T lymphocyte;
IFNγ: interferon-gamma;
IFNγ^hi^: IFNγ^high^ subset;
IFNγ^lo^: IFNγ^low^ subset;
WT: Wild-type;
GzmB-mTFP KI: Granzyme B-mTFP Knock-in;
TCR: T-cell receptor;
T_N_: Naive CD8+ T cells;
T_E_/T_EM_: Effector/effector memory T cells;
T_CM_: Central memory T cells;
T_EX_: T cell exhaustion;
CGs: Cytotoxic granules;
SGs: Secretory lysosomes;
CD44: Homing Cell Adhesion Molecule;
CD62L: L-selectin;
CD25: Interleukin-2 receptor;
CD127: IL-7 receptor alpha;
LAMPs: Lysosome-associated membrane glycoproteins; CD107a (LAMP-1), CD107b (LAMP-2), and CD63 (LAMP-3);
FasL/CD95L: Fas ligand;
Fas: Fas receptor;
CRTAM: Class I-restricted T cell-associated molecule;
DPBS: Dulbecco’s Phosphate-Buffered Saline;
BME: β-Mercaptoethanol;
IF: immunofluorescence;
ICC: immunocytochemistry;
FC: Flow cytometry;
TNFα: Tumor necrosis factor-alpha;
IL-6: Interleukin-6;
SIM: Structured illumination microscopy;
IRF-1: Interferon regulatory factor-1;
MFI: Median fluorescence intensity;
PCC: Pearson’s correlation coefficient.

## Acknowledgments

We thank Anja Bergsträßer, Margarete Klose, Nicole Rothgerber and Katrin Sandmeier for excellent technical assistance.

## Author Contributions

VP and XL conceived and designed the study. XL conducted flow cytometry experiments and performed data analysis. XL carried out structured illumination microscopy (SIM) imaging and completed co-localization analysis. JR, VP, and XL secured funding for the research. EK and H-FC provided critical insights for data interpretation. EK and VP offered technical support for flow cytometry and structured illumination microscopy. XL drafted the manuscript with input from all authors. All authors reviewed, edited, and approved the final version of the manuscript.

## Availability of data and materials

Additional data and information that support the findings of this study are available from the corresponding author upon reasonable request.

## Ethics approval and consent to participate

All experimental procedures were conducted in compliance with regulations of the state of Saarland (Landesamt für Verbraucherschutz, AZ.: 2.4.1.1 and 11/2021).

## Consent for publication

Not applicable.

## Additional information

**Extended Data Table 1.** Antibodies used in this study.

| Antibody, anti- | Supplier | Clone | Host | Dilution | Application |
| --- | --- | --- | --- | --- | --- |
| Mouse CD3e | BD Pharmingen | 145-2C11 | Hamster | 1:100 | Plate/glass coating |
| Mouse IFNγ | Biolegend | XMG1.2 | Rat | 1:200 | FC, ICC |
| Mouse IFNγ-Alexa488 | Biolegend | XMG1.2 | Rat | 1:200 | FC, ICC |
| Mouse GzmB-Alexa647 | Biolegend | GB11 | Rat | 1:200 | FC, ICC |
| Mouse-CRTAM-PE | Biolegend | 11-5 | Rat | 1:200 | FC |
| Mouse-CD25-FITC | BD Pharmingen | 7D4 | Rat | 1:200 | FC |
| Mouse-CD44-PE | BD Bioscience | IM7 | Rat | 1:1200 | FC |
| Mouse-CD62L-APC | BD Bioscience | MEL-14 (RUO) | Rat | 1:1600 | FC |
| Cis Golgi | BD Bioscience | GM 130 | Mouse | 1:100 | ICC |
| Mouse-CD107a-PE | BD Bioscience | (1D4B) | Rat | 1:200 | FC |
| Mouse-CD69-APC | Thermo Scientific | H1.2F3 | Hamst | 1:100 | FC |
| Mouse-IgG-Alexa647 | Thermo Scientific | — | Goat | 1:1000 | ICC |
| Rat-IgG-Alexa647 | Thermo Scientific | — | Chicken | 1:800 | ICC |
| Mouse-IgG-Alexa568 | Thermo Scientific | — | Goat | 1:400 | ICC |

**Note:** FC: flow cytometry, ICC: immunocytochemistry.

## Extended Data

**Extended Data Figure 1.**
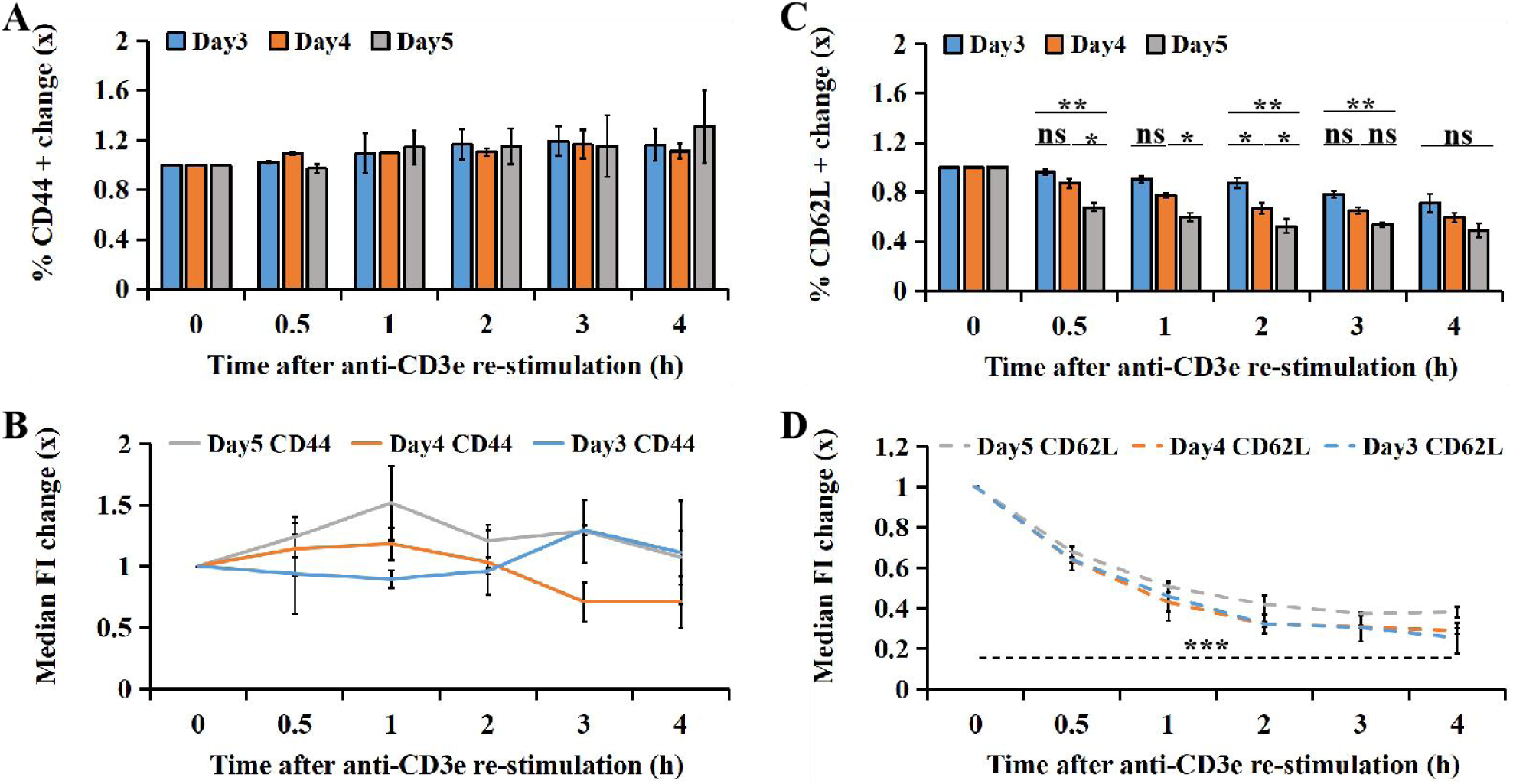
Dynamic expression profiles of CD44 and CD62L in reactivated CTLs. WT cytotoxic T lymphocytes (CTLs) activated for 3-5 days were re-stimulated with plate-bound anti-CD3e antibody (10 μg/mL) for the indicated durations (0, 0.5, 1, 2, 3, and 4 hours), followed by intracellular staining for IFNγ and granzyme B (GzmB). **(A, C)** Fold change in the percentage of CD44+ and CD62L+ cells. (**B,D)** Median fluorescence intensity (MFI) of CD44-PE (rat anti-mouse) and CD62L-APC (rat anti-mouse). Data represent mean ± SEM from ≥3 independent experiments. Statistical significance was determined by one-way ANOVA with post hoc unpaired t-tests: NS, *p* > 0.05; *0.01 < *p* ≤ 0.05; **0.001 < *p* ≤ 0.01; ***0.0001 < *p* ≤ 0.001.

**Extended Data Figure 2.**
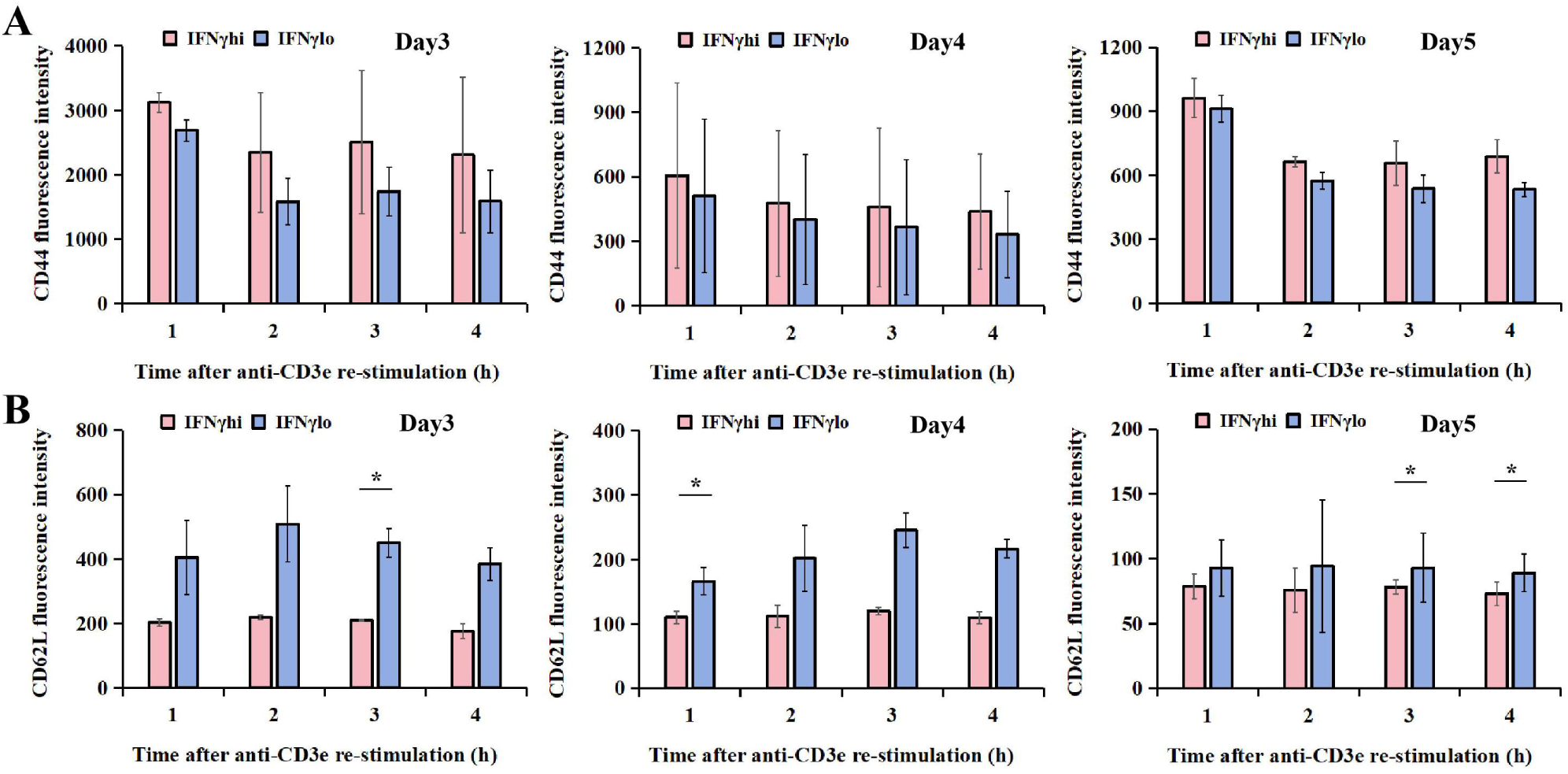
Flow cytometric analysis of T cell activation markers. WT cytotoxic T lymphocytes (CTLs) activated for 3–5 days were re-stimulated with plate-bound anti-CD3e antibody (10 μg/mL) for 0–4 hours and stained for surface expression of CD44 and CD62L. (**A)** Median fluorescence intensity (MFI) of CD44-PE (rat anti-mouse) in CD44+ IFNγ^hi^ and IFNγ^lo^ CTLs. (**B)** MFI of CD62L-APC (rat anti-mouse) in CD62L+ IFNγ^hi^ and IFNγ^lo^ CTLs. Data represent mean ± SEM from ≥3 independent experiments. Statistical analysis was performed using one-way ANOVA followed by unpaired t-tests: NS, *p* > 0.05; *0.01 < *p* ≤ 0.05; **0.001 < *p* ≤ 0.01; ***0.0001 < *p* ≤ 0.001.

**Extended Data Figure 3.**
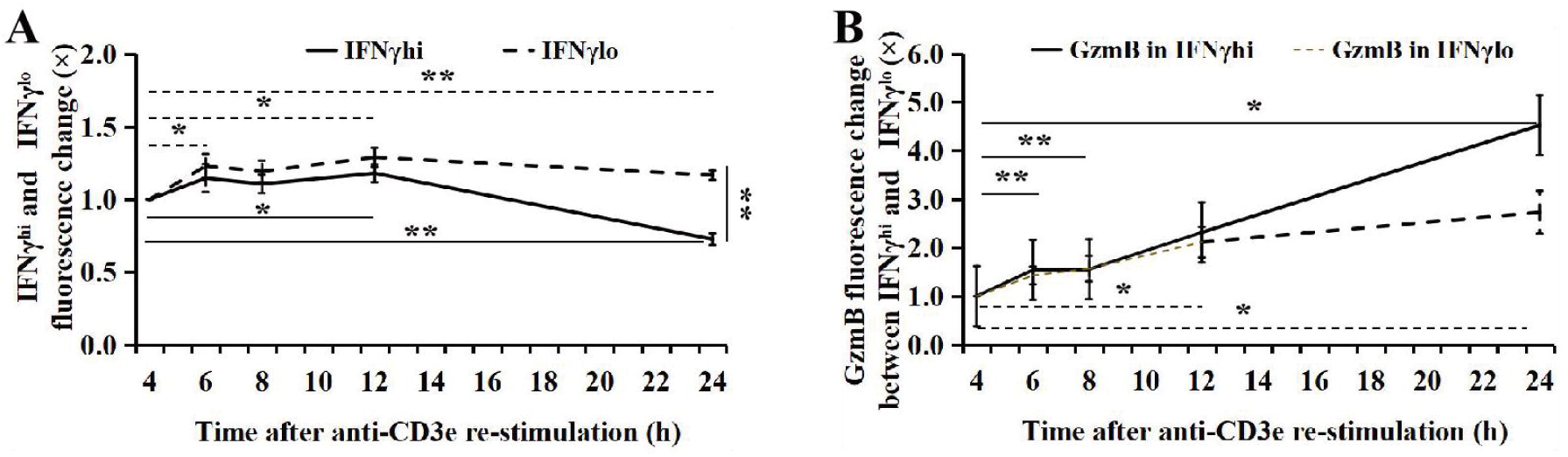
Temporal dynamics of IFNγ and granzyme B expression in activated CTLs. WT cytotoxic T lymphocytes (CTLs) activated for 4 days were re-stimulated with plate-bound anti-CD3e monoclonal antibody (10 μg/mL) for 4–24 hours. Cells were subsequently stained for intracellular IFNγ and GzmB using fluorescently labeled antibodies and analyzed by flow cytometry (FlowJo v10.10.0). (**A)** IFNγ fluorescence intensity changes in IFNγ^hi^ and IFNγ^lo^ subsets. (**B)** GzmB fluorescence intensity changes in IFNγ^hi^ and IFNγ^lo^ subsets. Data represent mean ± SEM from 3 independent experiments. Statistical significance was determined by one-way ANOVA followed by unpaired t-tests: NS, *p* > 0.05, *0.01 < *p* ≤ 0.05, **0.001 < *p* ≤ 0.01, ***0.0001 < *p* ≤ 0.001.

**Extended Data Figure 4.**
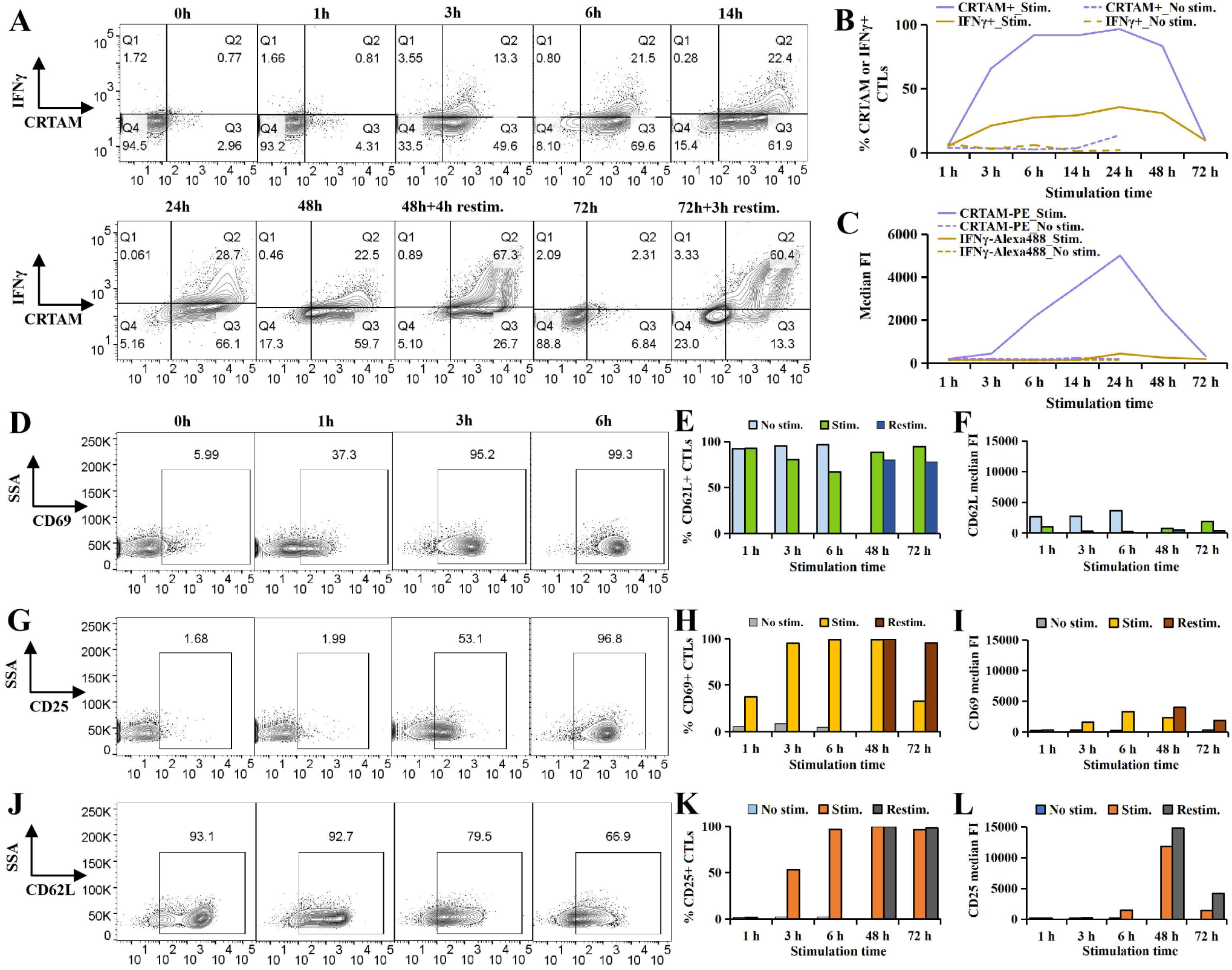
Temporal profiling of activation markers and IFNγ expression in primary CTLs. Day 0 WT cytotoxic T lymphocytes (CTLs) were stimulated in 96-well plates with plate-bound anti-CD3e (10 μg/mL) and anti-CD28 (5 μg/mL) antibodies for 0 h, 3 h, 6 h, 14 h, 24 h, 48 h,and 72 h, followed by surface staining for CRTAM, CD62L, CD69, and CD25, and intracellular staining for IFNγ. **(A)** Pseudocolor plots showing temporal expression patterns of CRTAM and IFNγ in day 0 CTLs following stimulation. (**D, G, J)** Pseudocolor plots depicting dynamic expression of CD62L, CD69, and CD25 in day 0 CTLs. **(B, E, H, K)** Quantification of CRTAM+, IFNγ+, CD62L+, CD69+, and CD25+ cell populations in stimulated day 0 CTLs. **(C, F, I, L)** Quantification of CRTAM-PE, IFNγ-Alexa488, CD62L-APC, CD69-APC, and CD25-PE median fluorescence intensity. Data represent mean ± SEM.

